# Self-assembly Controls Self-cleavage of HHR from ASBVd(-): a Combined SANS and Modeling Study

**DOI:** 10.1101/045054

**Authors:** Fabrice Leclerc, Giuseppe Zaccai, Jacques Vergne, Martina Řìhovà, Anne Martel, Marie-Christine Maurel

## Abstract

In the Avocado Sunblotch Viroid (ASBVd: 249-nt) from the Avsunviroidae family, a symmetric rolling-circle replication operates through an autocatalytic mechanism mediated by hammerhead ribozymes (HHR) embedded in both polarity strands. The concatenated multimeric ASBVd (+) and ASBVd (-) RNAs thus generated are processed by cleavage to unit-length where ASBVd (-) self-cleaves with more efficiency. Absolute scale small angle neutron scattering (SANS) revealed a temperature-dependent dimer association in both ASBVd (-) and its derived 79-nt HHR (-). A joint thermodynamic analysis of SANS and catalytic data indicates the rate-determining step corresponds to the dimer/monomer transition. 2D and 3D models of monomeric and dimeric HHR (-) suggest that the inter-molecular contacts stabilizing the dimer (between HI and HII domains) compete with the intra-molecular ones stabilizing the active conformation of the full-length HHR required for an efficient self-cleavage. Similar competing intra- and inter-molecular contacts are proposed in ASBVd (-) though with a remoter region from an extension of the HI domain.

## Introduction

Viroids, which may represent living vestiges of pre-cellular evolution in the hypothesis of an RNA World are the smallest and simplest replicating molecules known [1]. They consist of a single-stranded circular RNA genome, 246 to 375 nu-cleotides long; the genome does not code for proteins and replicates in plant cells. Avocado Sunblotch Viroid (ASBVd) is a member of the Avsunviroidae, which replicate in chloro-plasts; the viroid circular RNA (’positive’ polarity) is copied to a linear oligomer strand (’negative’ polarity) including a hammerhead ribozyme (HHR) sequence that cleaves the resulting strand at each place where the genome begins a repetition, yielding multiple copies. The monomeric fragments then reassume a circular shape, through ribozyme ends junction, to produce progeny viroid RNA [2, 3, 4]. Viroid plus and minus strands are not found in the same amounts in plant cells [5]. ASBVd (+) and ASBVd (-) display different folded structures, with a less stable (-) strand [6, 7]. ASBVd (+) may also adopt alternative conformations; furthermore, the (+) and (-) strands of the viroid are not auto-cleaved with the same efficiency, the activity of the ASBVd (-) having faster kinetics [8]. In a non-conventional host, ASBVd (+) and ASBVd (-) are not equally sensitive to exonuclease degradation [9].

Absolute scale small angle neutron scattering (SANS) is a highly appropriate method to determine structures and interactions in solution at low resolution, with complementary advantages over SAXS (small angle X-ray scattering) [10]: there is no radiation damage with neutrons; the RNA solution is in an easily accessible quartz cell and the identical sample can be measured in real time, while cycling over a range of temperatures. By using a Guinier analysis, two parameters were obtained and analysed on an absolute scale [11]: (i) the forward scattered intensity, from which was calculated the molar mass of the effective scattering particle to determine if it were the RNA monomer, dimer, or a mixture; (ii) the radius of gyration (in Å units), which is a measure of conformation. In a given salt condition and temperature, the analysis informs on whether the monomer is compact or extended and if the dimer is composed of a side-side or end-to-end association. For long particles, in which one dimension is much larger than the other two, a ‘long rod’ approximation analysis permits to derive the cross-sectional radius of gyration and mass per unit length, both on an absolute scale (in Å and gram/ Å units, respectively).

RNA molecules often include common catalytic motifs such as HHR, which can independently assume various activities either within a large molecule or excised from it. Viroids in nature are exposed to diverse environmental factors, such as circadian temperature variations, which impact the replication cycle [12]. In this paper, we report on a small angle neutron scattering (SANS) and molecular modelling study of ASBVd (-) and its excised HHR (-) as a function of temperature, which revealed a shared plasticity in behaviour related to catalytic function. Reversible temperature dependent dimer to monomer dissociation was observed for both HHR (-) and ASBVd (-). The structural results are correlated with an Arrhenius of catalytic activity in the same temperature range. The change in activation energy previously observed for catalytic activity at about 25 °C for a hammerhead type ribozyme [13] and by us for HHR (-) from ASBVd (-), from activity studies [14], is paralleled by a similar activation energy change for dimer to monomer dissociation. The stable end-to-end HHR dimer observed below the activation energy transition is poorly active. A joint thermodynamic analysis of SANS and catalytic activity data revealed a strong correlation between the dimer/monomer transition and the rate determining step of the reaction.

Since the ASBVd sequence is not symmetrical, the (+) and (-) strands are not equivalent and their respective catalytic activity also differs. ASBVd (+) can be found in different concatenated multimeric forms up to octamers while ASBVd (-) is just present as a monomeric or dimeric form because of a more efficient cleavage activity [15, 7]. This difference in activity can be attributed in part to the weak stability of the HHR (+) fold due to an unstable stem HIII (only two base-pairs) in the canonical HHR motif which is stabilized by one additional base-pair in the HHR (-) fold. The presence of tertiary contacts was also shown to have a direct impact on the catalytic activity of artificially ASBVd-derived hammerhead ribozymes [16]. These contacts, established between the two RNA domains (stem HI and coaxial stems HII and HIII), were proposed to play a role in the stabilization of the active conformation: mutants designed to remove those contacts exhibit reduced activities. More recently, it was shown that the presence of a single tertiary contact is sufficient to enhance the catalytic activity of a minimal hammerhead ribozyme [17].

In this study, we examine the role of the possible folded structures and tertiary contacts in the modulation of the dimer association/dissociation of HHR (-) and ASBVd (-) and we propose 2D and 3D models supported by the SANS data.

## 1 Materials and Methods

### Sample Preparation

Viroid And Ribozyme RNA Samples Asbvd And Hhr RNA Were Synthesized By In Vitro Transcription Of Dna Templates With T7 RNA Polymerase. For Asbvd, Dna Template Se-Quence From Plasmid Pbmasbvd-Hhr Was Amplified By Pcr With Taq Dna Polymerase Using Dna Primers, T7-Promoter Primer Taatacgactcactataggaagagattg-Aagacgagtg, And Reverse Primer Gatcacttcgtctctt-Cagg. For Hhr, Synthetic Dna Template (Commercial Prod-Uct From Eurofins Mwg Operon) Ggttcttcccatctttcc-Ctgaagagacgaagcaagtcgaaactcagagtcgga-Aagtcggaacagacctggtttcgtc, Was Pcr Ampli-Fied With T7-Promoter Primer Taatacgactcactatag-Gttcttcccatctttccctg And Reverse Primer Gac-Gaaaccaggtctgttccg. RNA Transciption Product For Asbvd Is 249 Nt Long (Two Supplementary G Nt Are Added At The 5’ End Of Natural Asbvd Sequence For Efficient Tran-Scription By T7 RNA Polymerase), And Hhr Is 79 Nt Long (Designed From Asbvd Hammerhead Ribozyme Plus One Extra G Nt At 5’ End For Efficient Transcription). After Transcription, Dna Template Was Degraded By Dnase I Treatment And RNA Was Purified By Electrophoresis On Denaturing (7 M Urea) 10 % Polyacrylamide Gel. Piece Of Gel Containing Full-Length RNA, Visualized By Uv Shadowing, Was Cut And RNA Eluted By Diffusion In 0.3 M Sodium Acetate, Quantified By Uv AbsorpTion At 260 Nm, And Ethanol Precipitated. After Centrifugation, RNA Pellet Was Suspended In Water And Stored At −20 °C.

### Absolute Scale Small Angle Neutron Scattering (SANS)

Ribozyme and viroid solutions at about 5 mg/ml in standard buffer (10 mM cacodylate pH 7.5 150 mM KCl) were examined on the D22 SANS camera at the Institut Laue Langevin, Grenoble (http://bit.ly/Institut-Laue-Langevin). Sample intensities were recorded with neutrons of wavelength l = 6 Å (10 % Δλ/λ). The scattering data were corrected for solvent background, efficiency of the detector cells, then radially averaged around the direct beam center and calibrated in absolute units by the scattering of 1.00 mm of H2O. The scattered neutron intensity was determined as the macroscopic cross-section I(Q) in units of [*cm*^−1^] versus the momentum transfer Q = (4*π*/λ)*sin*θ[Å^−1^] where 2q is the scattering an-gle. The sample-to-detector and collimation distances were both 5.6 m, providing a Q range of 0.008 < Q (Å^−1^) < 0.100. Sample volumes of about 150 micro-litres were contained in 1.00 mm path-length quartz cells. Temperature was controlled to within 1° by circulating water from a bath thermostat.

The radius of gyration of scattering contrast, R*_g_*, of a sample was determined from the scattered intensity, I(Q) by applying the Guinier approximation [11].

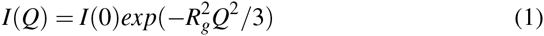

The molar mass (M) for each sample was extracted from the absolute scale value of I(0) by using equation (2) [11].

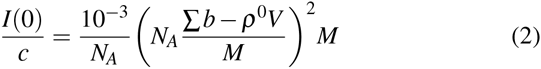

The factor 10^−3^ is required in order to express RNA concentration c (determined from OD at 260 nm) in mg/ml, N_A_ is Avogadro’s umber, the term in brackets is the excess scattering length of the RNA (in units of cm) per unit molar mass (Σ^b^ is the scattering length sum of RNA atoms, calculated from its composition; p is scattering length density of the solvent (cm^−2^); V is RNA volume calculated from its partial specific volume) and M is RNA molar mass in grams per mole.

In the case of a rod-like particle in which one dimension is significantly larger than the other equation (3) is a good approximation that yields the cross-sectional radius of gyration, R_*c*_ [18].

If the sample is poly-disperse, the I(0) and 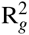 values rep-resent number averages:

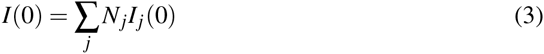

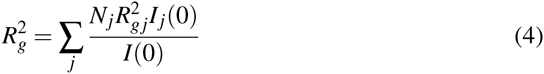

where N*_j_* is the number of particles j in the solution with radius of gyration R*_gj_*, and forward scattered intensity value I_*j*_(0).

Similarly to the mechanical radius of gyration of a body being equal to the distribution of mass (contrast in the case of scattering experiments) weighted by the square of its distance to the centre of mass, the cross sectional radius of gyration corresponds to the distribution in the cross section of a long rod projected on a two dimensional plane perpendicular to the axis of the rod. The radius of gyration of a long body is related to its cross-sectional radius of gyration and length L by equation (5):

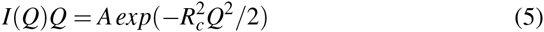

The molar mass per unit length (M/L) of the rod in gram/Å is calculated from A, the value of the function extrapolated to Q=0, by using equation (4)

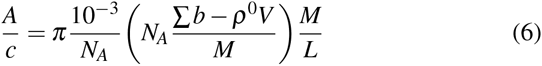

### Arrhenius analysis

In the Arrhenius equation,

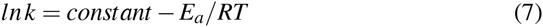

k is a rate constant, E_*a*_ is activation energy, R is the universal gas constant and T is temperature in degrees Kelvin. Activation energy is calculated from the slope of the straight line of the Arrhenius plot, ln k versus 1/T. Arrhenius plots were calculated from the rate of dimer dissociation of HHR (-)/Mg revealed by SANS (the amount of dissociation after half an hour at the given temperature), and from catalytic rates for ASBVd (-)/Mg activity published previously [14].

## RNA 2D Modeling

The Possible Modes Of Interaction Between Two Monomers Were Calculated Using Intarna [19] Using A Seed Sequence Of 7 Residues. The Web Server Version (Http://Bit.Ly/Intarna Was Used To Submit The Calculations Using The Specified Seed Size And The Three Different Temperatures At Which The Experi-Mental Measures Were Carried Out (10 °C, 25 °C And 45 °C). The Results Of The Calculations For HHR (-) And Asbvd (-) Are Provided In A ZIP Archive As Supplementary Material (Supple-Mentary Data). Intarna Was Previously Tested On The PAL2 Sequence From A Gamma Retrovirus (79-Nt RNA) To Reproduce The RNA-RNA Interaction Of A Metastable Dimer [20]. In The 2D Structure Corresponding To The 3D Dimer Model, The Interaction Energies Were Calculated Using RNAeval (Vienna Package [21]).

## RNA 3D Modeling

The 3D structure of the full-length hammerhead ribozyme from *Schistosoma mansoni* [22] (PDB ID: 2OEU [23]) was used as template to generate the 3D models of the monomer. The MMTSB toolkit [24] was used to make the mutations corresponding to the sequence of the hammerhead ribozyme from ASBVd (-). The paired interface between the two monomers involving a terminal loop including seven residues was taken from the 3D structure of the adenosylcobalamin riboswitch (PDB ID: 4GMA [25]; residues 75 to 83 and 191 to 203). The terminal loop and the 3’ end region of the two interacting monomers were modified and assembled based on the 3D coordinates of this interface. The 3D structure of the dimer model was optimized using the CHARMM program (minimization performed using a tolerance criterion of 10^−1^ kcal/mol Å) and the CHARMM27 forcefield [26] using modified parameters for the non-bonded interactions corresponding to an implicit solvent model [27]. The 3D model of the unbound conformation is very similar to the X-ray structure used as template (Fig. S1A). The bound conformations of each monomer are slightly remodeled in terminal loop of the HII domain and 3’ end region of the monomers 1 and 2, respectively (Fig. S1B-C).

## 2 Results

HHR (-) and ASBVd (-) in standard buffer (see Methods) and with added 2 mM Mg**^++^** (samples HHR (-)/Mg and ASBVd (-)/Mg) were examined by SANS as a function of temperature. At the first temperature point, the HHR(-)/Mg samples were measured before in-beam adjusting of the solution to 2 mM Mg**^++^** to confirm that the addition of Mg**^++^** did not modify the scattering curve significantly. The quality of the SANS data is illustrated in Figure 1 for HHR (-)/Mg at two temperatures. Similar quality data were obtained for all samples and conditions.

**Figure 1:**
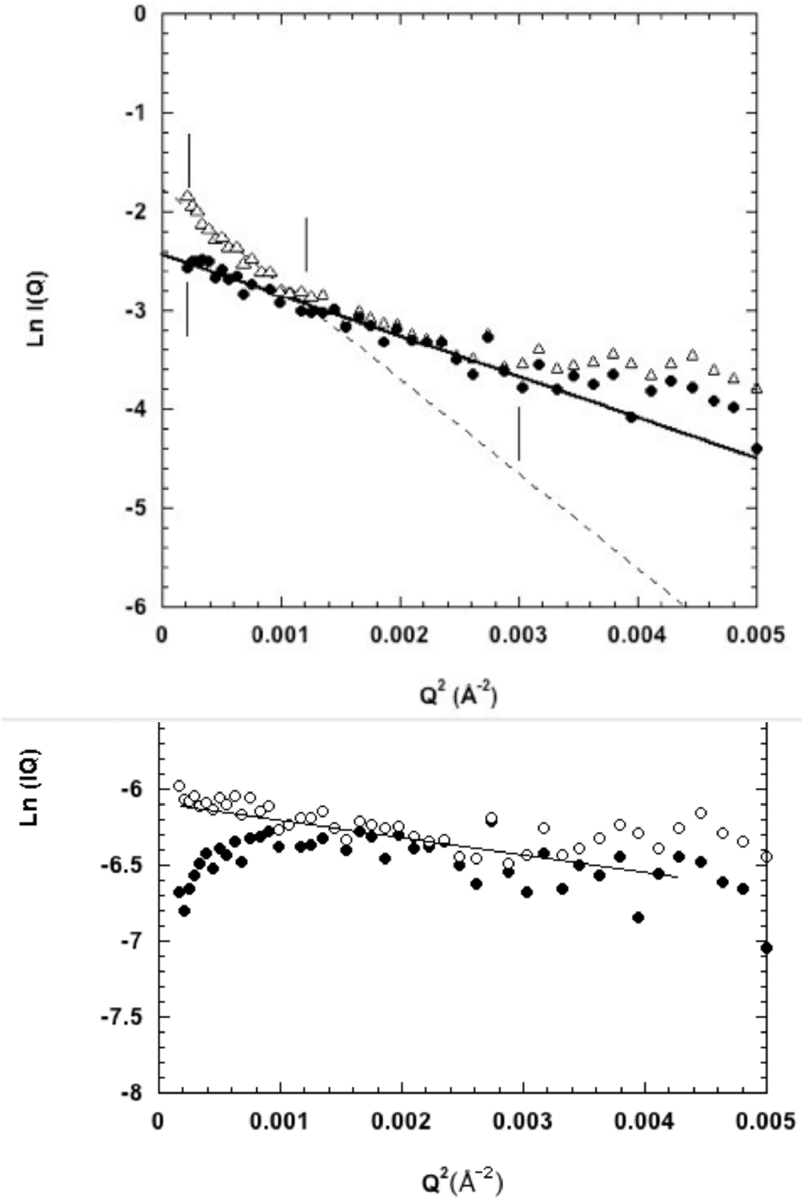
SANS Guinier plots of HHR (-)/Mg. (Top) Plot at low temperature (open triangles) and high temperature (filled circles). (Bottom) Plot in the rod representation (see Methods).

Scattering intensities *I* (*Q*) (*Q* **=** (*4*π*sin*θ)*/ λ*) where *2*θ is scattering angle and *l* is incident wavelength) were analysed according to the Guinier approximation (see Methods) to obtain on an absolute scale: radius of gyration (*R_g_*) in Å units and forward scattered intensity (*I*(*0*) in units of cm^−1^. The molecular mass of the effective scattering particle was calculated from the *I*(*0*), the concentration and the calculated scattering length density (SLD) contrast of HHR (-) and ASBVd (-) RNAs (see Methods). For each sample, measurements were taken at fixed temperatures in the following sequence: 6 °C, 10 °C, 25 °C, 45 °C, 25 °C, 10 °C, 25 °C, 45 °C, 10 °C.

The *I*(*0*) values of the ASBVd(-) samples plotted against temperature in Figure 2 show a reversible low level of particle association at low temperature (e.g. accounted for by 5% of the particles forming dimers) with full dissociation to monomers at the higher temperature, both in presence and absence of Mg**^++^**.

**Figure 2:**
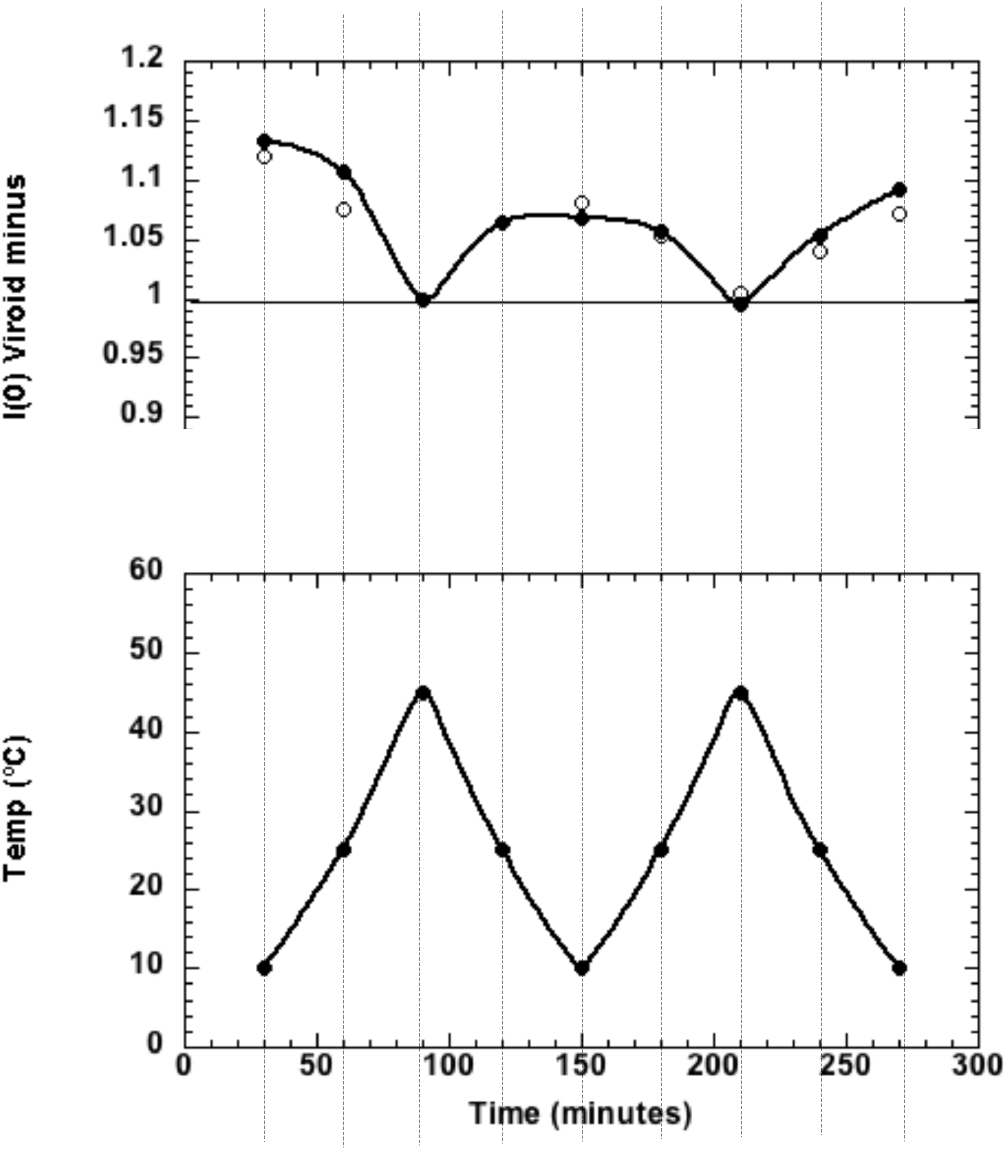
Forward scattered SANS intensity as a function of temperature for ASBVd (-). (Top) In absence (filled circles) and in presence of Mg**^++^** (open circles), at the corresponding temperature given in the (Bottom) plot. Exposure time at each temperature was 30 mins. The value at 1.00 corresponds to the intensity expected from the particle monomer.

Corresponding data for HHR are shown in Figure 3. The M, D lines correspond to the values expected for the HHR monomer and dimer respectively. HHR (-) particles in absence of magnesium ions start as dimers of *R_g_* ***~*** 50 Å at low temperature and fully dissociate to monomers of *R_g_* ***~*** 31 Å at 45 °C. In the following cooling cycle re-association was very small with the monomer still dominating the scattering. With added 2 mM Mg^++^, HHR (-)/Mg strands displays a striking temperature driven oscillatory behaviour as a function of temperature, between dimer-dominated scattering at low temperature and monomer-dominated scattering at the higher temperature.

**Figure 3:**
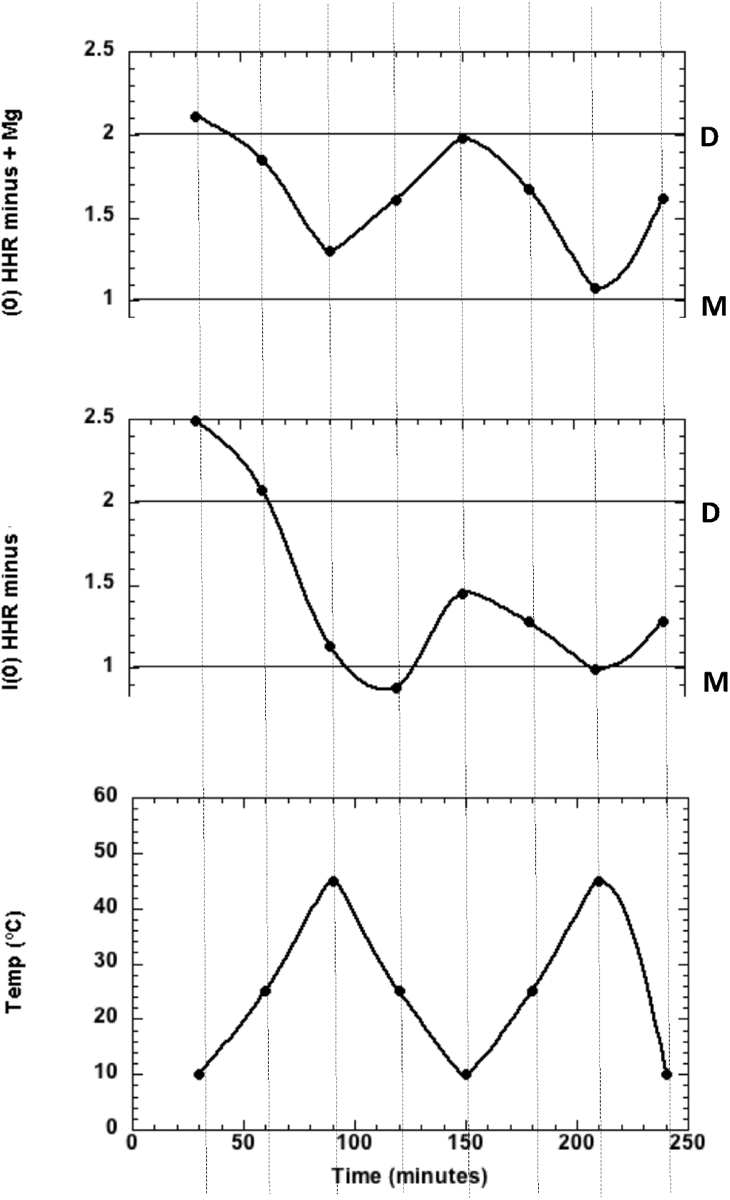
Forward scattered SANS intensity for HHR (-). (Top) In presence of Mg^++^, (Middle) In absence of Mg^++^, at the corresponding temperature given in the (Bottom) plot. Exposure time at each temperature was 30 mins. The values at 1.00 and 2.00 correspond to the intensity expected for the particle monomer (M) and dimer (D), respectively.

The data plotted according to the ‘long rod’ approximation (see Methods and Figure 1) indicate that both monomer and dimer are elongated particles with the same cross-sectional radius of gyration *R_c_* ***~*** 15 **±** 3 Å slightly larger than a RNA A helix and a mass per unit length of ***~*** 255 g/ Å very close to the value expected for a RNA A helix (one base pair per 2.3 Å is: 276 g/ Å). The length of the monomer molecule was calculated from the radius of gyration and cross-sectional radius of gyration values (see Methods) to be 96 Å. In the 3D model of HHR(-) generated by homology modeling (see Methods), the maximum distance is 96.7 Å (between the O2 and O1P atoms of residues 45 and 77, respectively). Note that the length of a 40 base pair RNA A helix is 92 Å. The dimer *R_g_* of 50 Å fits approximately with an end-to-end association of two monomers.

The RNA 3D model is consistent with the canonical HHR fold and maintains the tertiary contacts that stabilize the active conformation as indicated in the diagram of the corresponding 2D RNA structure (Fig. 4A). The optimal mode of interaction (as calculated by IntaRNA: see Methods and Supplementary Material) involves the pairing between the regions 72-79 and 26-33 of both monomers (seed regions: 72-78 and 27-33): only stem HII from the second monomer is unfolded (Fig. 4B). The 3D model of the dimer was built using slightly different seed regions: 70-76 and 29-35 derived from the sub-optimal mode of association and extended in the terminal loop from stem HII (Fig. 4C). This mode of association preserves the native 2D folds of both monomers (Fig. 5A). However, all the tertiary contacts are removed in the second monomer because the nucleotides 29, 31 and 34 are part of the seed region. The hybridization requires a remodeling of the terminal loop from stem HII (Fig. 5A-B). In the dimer (Fig. 5C), only the first monomer retains both the native 2D and 3D folds while the second monomer loses the tertiary contacts that stabilize the active conformation. For more details, a summary view is provided that compares the 3D structure of a minimal HHR (Fig. S1A) with the modeled 3D structures of both monomers (Fig. S1B-C); a focus on the tertiary contacts is shown.

The active conformation of HHR (-) is more compact due to the tertiary contacts between the two domains. In the morphing simulation from the less-active to highly active conformation of HHR performed by O’Rourke *et al*. [17], the ratio between the radius of gyration of the two conformations is around 1.1 and can be considered as the minimum ratio since the calculation is done on a minimal HHR (where the deviation will probably be under-estimated compared to a full-length HHR). The 3D model proposed for the monomer corresponds to the active conformation of a typical full-length HHR; the calculated radius of gyration is: *R_g_* = 26 Å and 29 Å when corrected with the scaling factor of 1.1, which is pretty close to the expected value of 31 Å assuming the monomer is mostly in some inactive conformation in solution. The scaling factor between the inactive/active conformations can then be adjusted to 1.2 as calculated between the expected/calculated values for the monomer (31/26). For the dimer, the calculated radius of gyration is: *R_g_* = 35 Å; the corrected value would be 42 Å which is still significantly below the expected value of 50 Å indicating the 3D model over-estimates the compactness of the dimer. In the second monomer, the loss of all the tertiary contacts between the two RNA domains (Fig. S1 C) would presumably make it much more flexible than the inactive conformation of the free monomer. The particular flexibility of the second mononer is thus expected to increase the radius of gyration of the dimer (the scaling factor would then be 1.4 instead of 1.2). The local unfolding of stem HII from the second monomer could increase even more the radius of gyration to optimize the inter-molecular interaction (Fig. 4). Recently, the active conformation of a WT HHR was determined by X-ray crystallography (PDB IDs: 5DI2, 5DI4, 5DQK) in a dimeric form where the two monomers are associated through the HI domain [28]. The radii of gyration calculated for this intermolecular dimer on a WT HHR of the same length (79-nt) indicates a more compact mode of association by 3 to 4 Å (Fig. S2).

The hybridization energy calculated for the seed region of the 3D model gives the following trend: −9.3 kcal/mol at 10 °C, −8.5 kcal/mol at 25 °C and −5.8 kcal/mol at 45 °C (Fig. 5). At this latter temperature, the dimer is mostly dissociated in two monomers which can both adopt an active conformation only limited by the dynamics of the monomer thus enhancing the global catalytic activity.

In this study, we consider a folded form of the ASBVd (-) RNA which is compatible with the HHR (-) fold and catalytic activity (Fig. 6A). In this two hairpin loops structure, the 5’ end of the HHR motif is located just after the first hairpin loop and its 3’ end closes the second hairpin loop at the 3’ terminal end of the RNA (Fig. 6A). The optimal interaction for the dimer requires a local unfolding of the two regions corresponding to the seed regions of association (Fig. 6B). The first region overlaps with an internal loop located at positions 137-139/178-185 and would open up two base-pairs upstream (136-137/185-186) and four base-pairs downstream (139-142/178-181). The second region corresponds to the three last base-pairs of the second hairpin loop that also closes the HII stem of the HHR motif (66-68/244-246). The interaction involves the seed regions 136-142 and 241-247 making the residues 241 and 243 (equivalent to the positions 29 and 31 in HHR (-), Fig. 5A) unavailable for the tertiary contacts that are normally established with the respective residues 224 and 222 (equivalent to the positions 12 and 10 in HHR (-), Fig. 5A). A sub-optimal self-association mode (ΔΔ*E* ∼ 2 kcal/mol) also involves a possible inter-molecular contact with the position 222 (Supplementary Data).

In order to explore the functional relevance of the SANS observations, an Arrhenius plot was calculated from the rate of HHR (-)/Mg dimer dissociation as a function of temperature and compared to a similar plot calculated from catalytic data on the same ribozyme published previously [14]. The activity and rate of dissociation plots in Figure 7 are strikingly similar, showing a break at about 27 °C between regimes of activation energy 7 ± 1 kcal/mole and 25 ± 5 kcal/mol, at higher and lower temperatures, respectively.

## 3 Discussion

In HHR (-), the dimer dissociation is correlated with a better activity (Fig. 7). In our model, the dimer association of HHR (-) is based on the inter-molecular interaction between two folded HHR motifs. The monomer/dimer equilibrium profile at different temperatures suggests the association/dissociation between two monomers is reversible with a low energy barrier (Fig. 3). We can draw a parallel with the dimer formation at low temperature of a 79-nt RNA used to study the competition between inter-molecular and intra-molecular folding [20]. It was shown that the inter-molecular association is restrained to some regions which are mostly single-stranded and preserves the intra-molecular fold of the more stable hairpin structure.

In the proposed model, the inter-molecular contacts between the two HHR (-) monomers preserve the 2D native folds of each monomer but compete with the tertiary intra-molecular contacts which stabilize the active conformation. In the dimer, only one of the two monomers can adopt the active conformation through the native tertiary contacts. Thus, only half of the HHR (-) would be fully active in the dimeric form. Once the dimer is dissociated, the residues involved in the inter-molecular contacts are released and can stabilize the active conformation in both monomers. The 3D structure of an artificial HHR crystallized as a dimer was recently determined [28]; it suggests an alternative mode of association where the seed regions (6 nt) correspond to the basal part of the HI stem. However, the radius of gyration for this mode of association is smaller and thus less consistent with the SANS data (Fig. S2B-C). On the other hand, we would not expect any modulation of the catalytic activity between the monomeric and dimeric forms since the two active sites are symmetrical and equivalent.

The dimer/monomer ratio is relatively stable along the two cycles of temperatures in the presence of Mg^++^: from 10 °C to 45 °C (2.1 and 1.9 in the first cycle at 10 °C, Fig. 3 (Top)) but it tends to decrease in the second cycle (1.9 and 1.6 in the second cycle, Fig. 3 (Top)). In the absence of Mg^++^ (Fig. 3 (Middle)), the decrease is more pronounced suggesting the influence of the ionic strength and/or divalent metal ion on the equilibrium. Alternative conformations may be generated at 45 °C after the first cycle which prevent a full re-association of the monomers.

The optimal dimer interaction of the 249-nt ASBVd RNA requires a local unfolding of: (1) the base-pairs surrounding an internal loop (2 upstream and 4 downstream base-pairs), (2) the three terminal base-pairs from the stem HII of the HHR motif next to the internal loop of the 5’ end of the HHR motif. Both dimer models for HHR (-) (Fig. 4) and ASBVd (-) (Fig. 6) are based on a weak interaction between accessible or partially accessible regions of the folded monomers; the calculated interaction between the seed regions is equivalent between HHR (-) and ASBVd (-): −5.8 kcal/mol and −5.6 kcal/mol, respectively. The lower dimer/monomer ratio of ASBVd (-) at 10 °C (Fig. 2) compared to that of HHR(-) (Fig. 3) may be explained by a larger energy cost to unfold and make the seed regions fully accessible in ASBVd (-).

**Figure 4:**
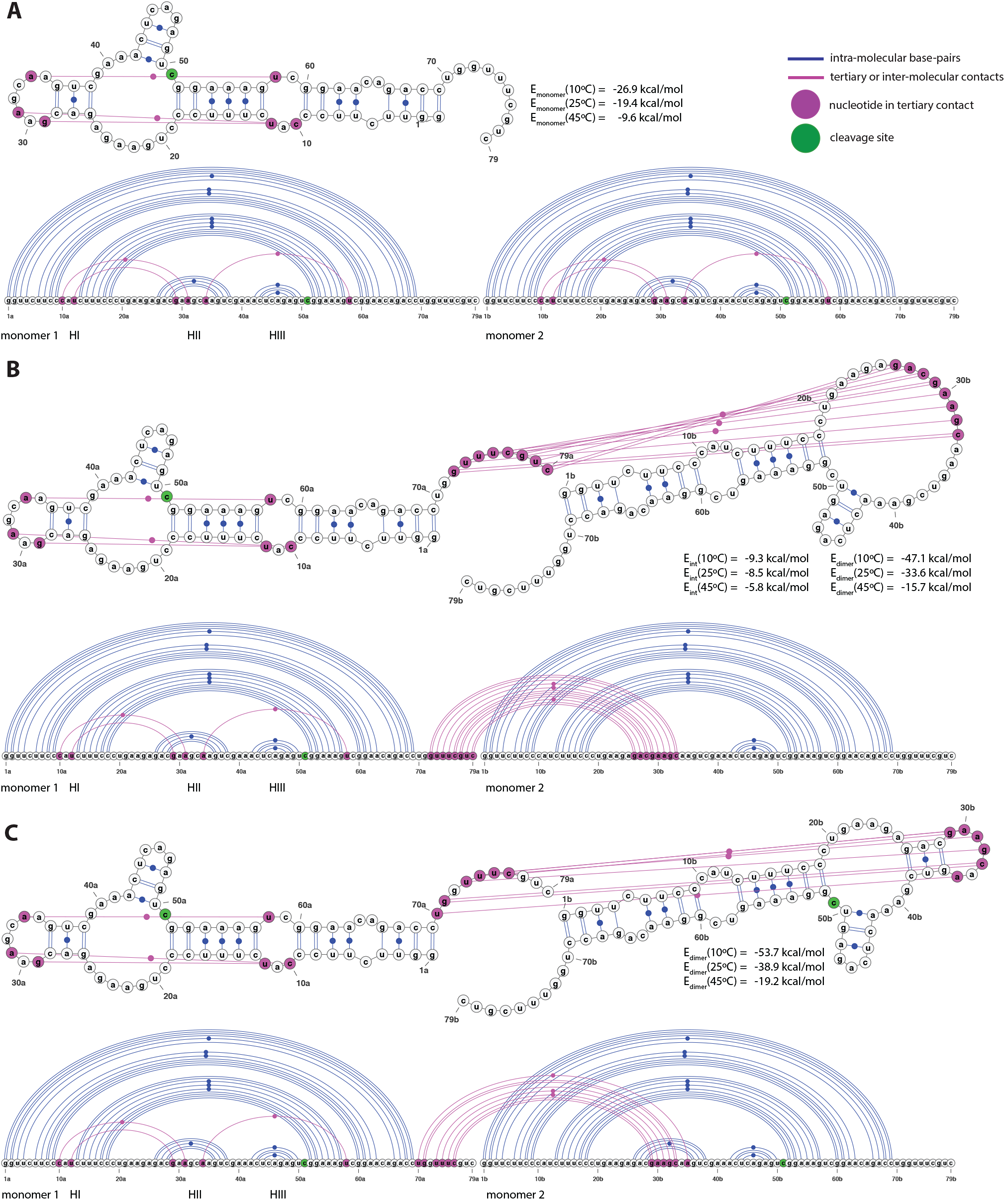
2D and linear representations of HHR(-). (A) Non-interacting monomers: only the intra-molecular base-pairs and tertiary contacts are indicated. (B) Optimal dimer hybrid: the inter-molecular interactions include the base-paring between the residues 72-79 from monomer 1 and 26-33 from monomer 2 (the seed regions correspond to 72-78 and 27-33). (C) Dimer hybrid used in the 3D modeling: only the base-pairs from the optimal dimer hybrid that preserve the native fold of both monomers are indicated. The interaction and hybrid energies are calculated for the pairing regions and for the full dimer at 10 °C, 25 °C and 45 °C using IntaRNA and RNAeval from the Vienna package, respectively (see Methods).

**Figure 5:**
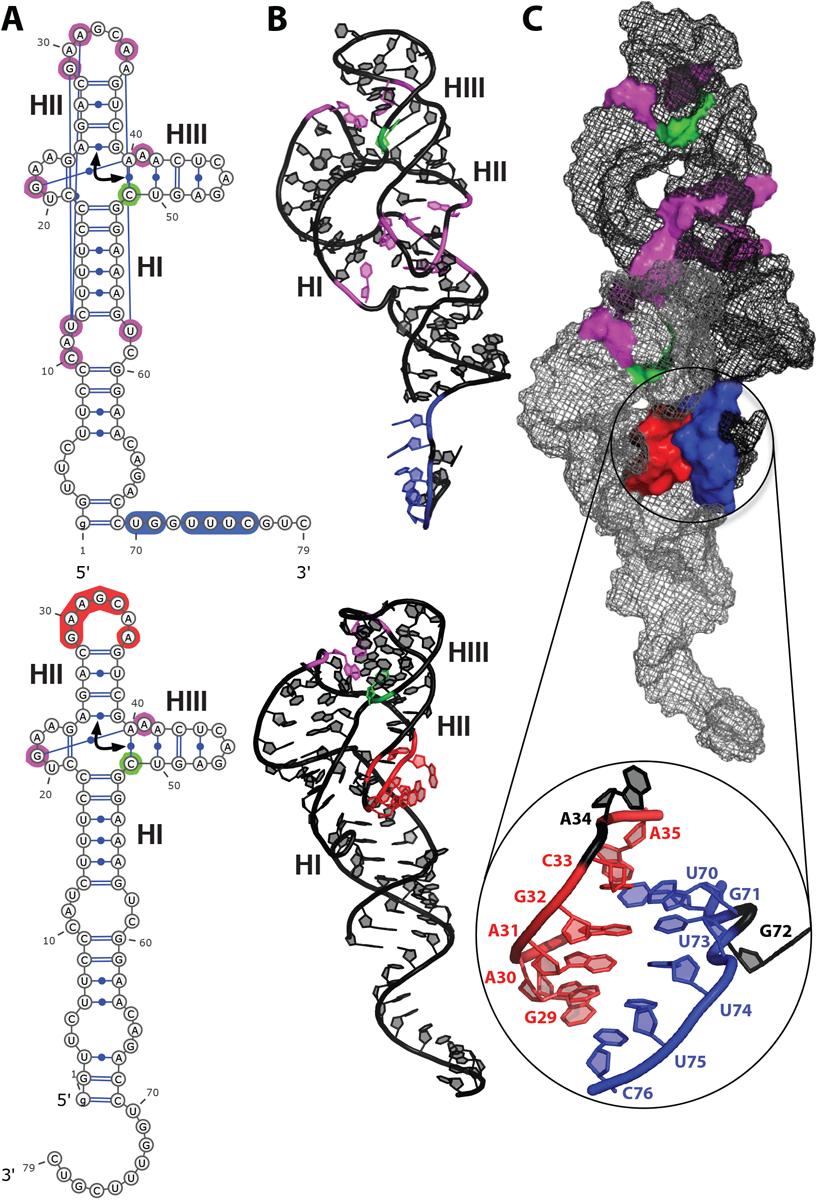
2D and 3D Models of HHR (-). (A) 2D structures of the monomer: intra-molecular tertiary contacts (magenta), inter-molecular paired regions (3’ end of HI domain: blue, terminal loop of HII domain: red), cleavage site (green), the arrow indicates the coaxial helices. (B) 3D modeled structures of the monomers shown in their respective orientations and bound conformations (inter-molecular contacts: blue and red; intra-molecular contacts: magenta; cleavage site: green). (C) 3D modeled structure of the dimer and its main interface: the two monomers are shown using a mesh representation (monomer 1: black; monomer 2: grey), the details of the interface is zoomed in on the seed regions between the two monomers (blue and red).

**Figure 6:**
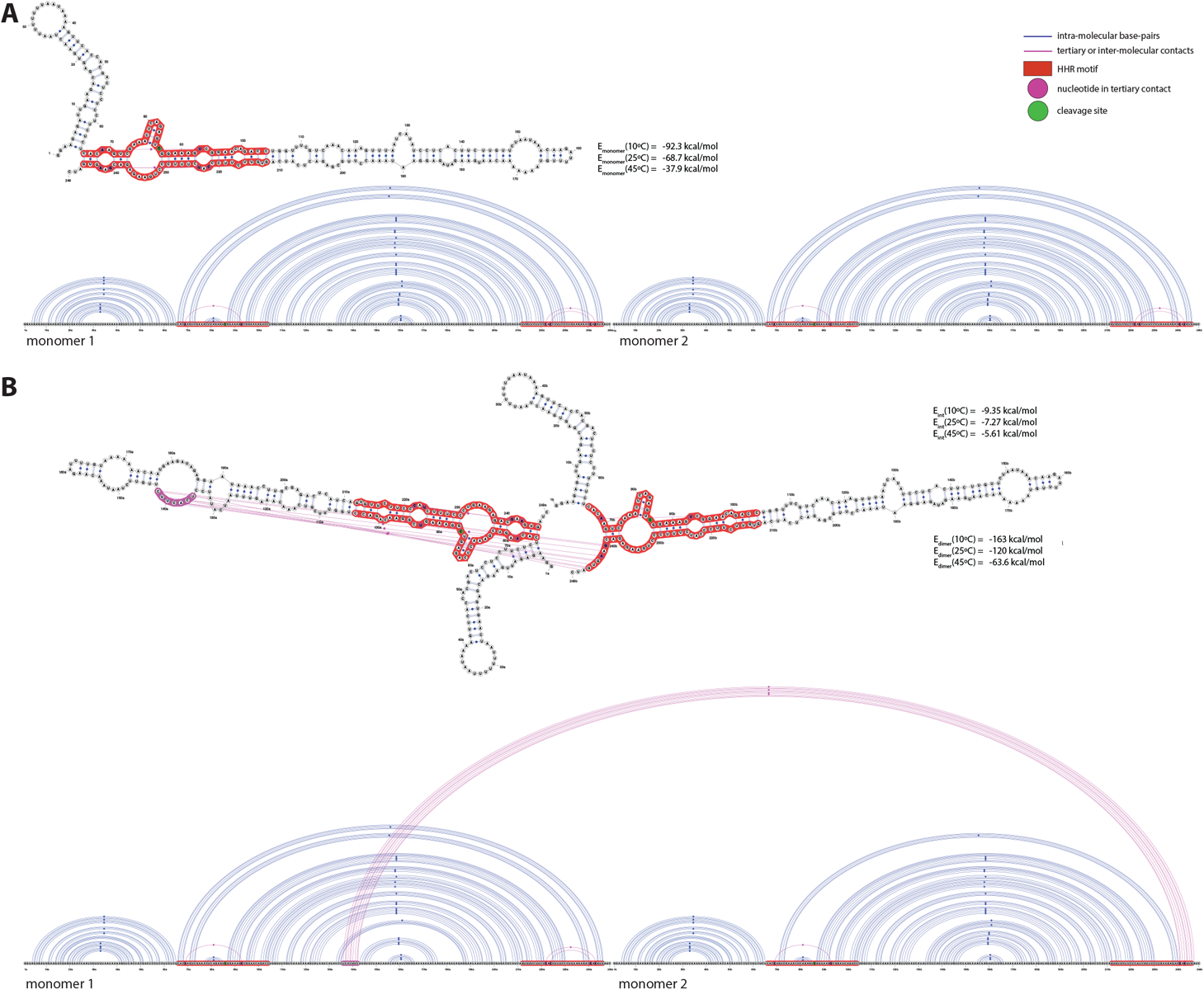
2D and linear representations of ASBVd (-). (A) Non-interacting monomers: only the intra-molecular base-pairs and tertiary contacts are indicated. (B) Optimal dimer hybrid: the inter-molecular interactions include the base-paring between the residues 136-142 from monomer 1 and 241-247 from monomer 2 (corresponding to the seed regions).

**Figure 7:**
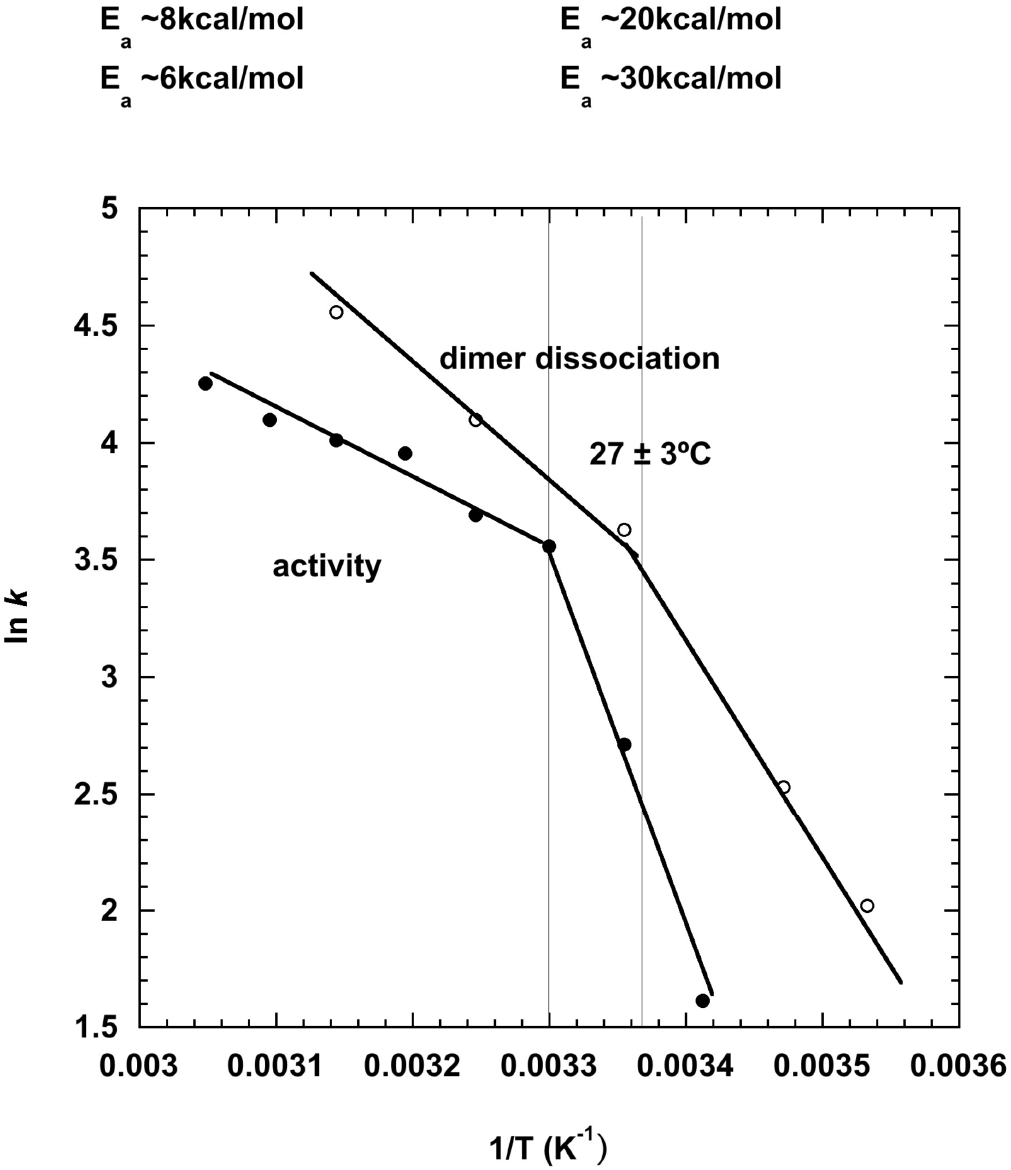
Arrhenius plots of HHR (-)/Mg rate of dimer dissociation as a function of temperature. The measures are those observed in SANS (open circles) and for HHR (-)/Mg catalytic activity (filled circles) (calculated from previous data [14]).

In vitro, the ASBVd (+) and ASBVd (-) RNAs behave in a different way with a 3.5 faster self-cleavage rate for ASBVd (-) [6]. As a result, the monomeric or cocatenated dimeric forms of ASBVd (-) prevail in vivo while ASBVd (+) is detected as various multimeric forms up to 8-mers [15, 7]. As proposed before, this rate difference can be explained by the ability to fold into a 2D structure compatible with a stable HHR motif (more stable in HHR (-) than in HHR (+)) but also by stabilizing the active conformation. The temperature is a key factor both in terms of folding and catalysis. In a range of temperatures between 20 **°**C and 60 °C, ASBVd (+) was shown to adopt multiple concurrent folds while ASBVd (-) seems to adopt a unique stable fold [8]. At 20 °C, the ASBVd (+) fold is more stable with a higher double helical content [6]. At 45 °C, the higher flexibility of ASBVd (-) is correlated with a better self-cleavage activity because of a HHR-compatible fold and a stabilization of the active conformation in presence of Mg^++^.

The in vitro transcripts ASBVd (+) and ASBVd (-) were shown to adopt a concatenated dimeric form corresponding to the so-called “double-HHR structure” which is the product of two monomeric HHR structures merged between the two HIII terminal loops [29, 30]. An alternative dimeric form also exists corresponding to two concatenated and also covalently linked monomers and it is referred as “single-HHR structure” [30]. In this particular structure, the two HHR motifs are merged between the two HI domains. It probably folds into a 3D structure similar to that of the dimer determined recently [28] where the 5’ and 3’ ends of both monomers would be ligated. Thus, the “single-HHR structure” would not allow any other inter-molecular contacts apart from those involved in the pairing of the HI stems because of distance constraints in the 3D structure (Fig. S2B-C). In this case, we should not expect any modulation of the catalytic activity between the monomeric and dimeric forms of ASBVd (-). During transcription, the “double-HHR structure” for both ASBVd (+) and ASBVd (-) were shown to be active. On the other hand, only ASBVd (-) can self-cleave as a “single-HHR structure”. Our dimeric model for ASBVd (-) is based on the association of two distinct molecules which can interact between each other through different remote seed regions (Fig. 6). For this model, we do expect a difference of activity between the monomeric and dimeric forms. The influence of the ASBVd (-) dimerization on the viroid replication and cycle in the plant remains to be evaluated.

## Acknowledgments

We are happy to acknowledge L. Porcar of ILL for help with the SANS instrumentation and data collection. We thank V. Lefort and L. da Silva for technical help. We are grateful to WG. Scott for providing the coordinates of the HHR structures generated in the morphing simulations.

